# A full-proteome, interaction-specific characterization of mutational hotspots across human cancers

**DOI:** 10.1101/2019.12.20.885293

**Authors:** Siwei Chen, Yuan Liu, Yingying Zhang, Shayne D. Wierbowski, Steven M. Lipkin, Xiaomu Wei, Haiyuan Yu

## Abstract

Rapid accumulation of cancer genomic data has led to the identification of an increasing number of mutational hotspots with uncharacterized significance. Here we present a biologically-informed computational framework that characterizes the functional relevance of all 1,107 published mutational hotspots identified in ∼25,000 tumor samples across 41 cancer types in the context of a human 3D interactome network, in which the interface of each interaction is mapped at residue resolution. Hotspots reside in network hub proteins and are enriched on protein interaction interfaces, suggesting that alteration of specific protein-protein interactions is critical for the oncogenicity of many hotspot mutations. Our framework enables, for the first time, systematic identification of specific protein interactions affected by hotspot mutations at the full proteome scale. Furthermore, by constructing a hotspot-affected network that connects all hotspot-affected interactions throughout the whole human interactome, we uncover genome-wide relationships among hotspots and implicate novel cancer proteins that do not harbor hotspot mutations themselves. Moreover, applying our network-based framework to specific cancer types identifies clinically significant hotspots that can be used for prognosis and therapy targets. Overall, we demonstrate that our framework bridges the gap between the statistical significance of mutational hotspots and their biological and clinical significance in human cancers.

## Introduction

Through DNA sequencing of tumor mutations, precision oncology has enabled the identification of cancer drivers, therapy targets and prognostic mutations that can guide individualized therapies for many cancer patients. For example, what was once defined as melanoma is now delineated as BRAF-positive or BRAF-negative melanoma, a meaningful distinction with respect to therapy with BRAF and MEK pathway inhibitors. Similarly, whether a tumor has deficient DNA mismatch repair defines whether the patient is eligible for immune checkpoint inhibitor monoclonal antibody therapy. Precision medicine now has become part of mainstream oncology and in 2019, >80% of oncology drugs in development are personalized medicines^1^. However, an important current limitation to precision medicine is the overwhelming number of total somatic mutations that accumulate during tumorigenesis and progression. A significant challenge is distinguishing bona fide driver mutations that promote tumor growth from passenger mutations that are neutral and have no mechanistic impact. To date, international efforts in cancer genomics have provided whole-exome sequencing for tens of thousands of human cancers^2-4^. Subsequent computational analyses have identified cancer driver genes in which mutations occur more frequently than expected^5-11^. Yet not all mutations on driver genes are driver mutations. This is usually interpreted as the driver-passenger paradigm, where the few recurrent mutations are viewed as drivers while most mutations, especially rare ones, are passengers that do not involve in oncogenesis^12,13^. In this regard, statistical models were developed to detect mutational hotspots (highly recurrently mutated residues across tumor samples) as candidate drivers. Such candidate list was quickly populated by over 1,000 hotspots^14,15^, but only a small number of them have well-defined functional consequences. It was recently reported that some hotspot mutations are in fact passengers that arose from the preference of APOBEC3A, a cytidine deaminase, for DNA stem-loops^16^. Thus, given the increasing number of cancer hotspots with uncertain significance, there is an urgent need to characterize their functional relevance towards translating the wealth of genomic data into biological and clinical insights.

Although it is now possible to systematically test certain mutations by experiments^17^, genome-wide prioritization of candidate driver mutations still involves bioinformatics tools that predict the impact of mutations on protein function at the individual protein level^18-20^. However, not all mutations are simple to interpret as causing a gross loss of protein. Many cancer mutations exert their oncogenic effects through altering specific aspects of protein activity and give cancer cells a selective advantage. One promising route to decipher this complexity is from the view that the cell is a network of interacting biomolecules where proteins carry out diverse functions by interacting with other proteins. We have previously demonstrated that one key feature in understanding the functional impact of mutations is whether they fall in the binding interfaces that mediate interactions with other proteins and critically, which specific interactions they mediate^21,22^. While studies of known disease mutations have already reported a strong association with protein interaction interfaces^23,24^, application of this feature has been largely limited by low coverage of structural information on interacting proteins; co-crystal structures and homology models together cover merely ∼6% of all known human interactions^25^.

In this study, we leverage our newly-established, the first human full-proteome 3D interactome with residue-resolution interface predictions (Interactome INSIDER^25^) to systematically identify protein-protein interactions that are affected by mutational hotspots, which constitutes a proteome-wide functional characterization of candidate driver mutations in cancer (**Fig. 1a**). Interactome INSIDER is a unified machine-learning framework that predicts partner-specific interaction interfaces for the entire experimentally determined human interactome, taking full advantage of all available 3D structural information for proteins and their interactions. By mapping mutational hotspots within this 3D interactome network, we analyze their impact not only at the individual protein level, as traditionally has been done, but extend beyond and interrogate their network properties to evaluate their oncogenic potential by how they may affect protein interactions. A unique advantage of our approach lies in its capacity to dissect specific interactions each hotspot affects while leaving others unchanged. Furthermore, instead of analyzing each hotspot individually as is commonly done in cancer driver mutation prediction^26-29^, we connect the effects of all hotspots to construct a hotspot-affected interactome network at the full proteome scale. Such network has not been possible to construct in any previous studies since interactome-based approaches suffered from either low coverage (i.e., focusing on only interactions with cocrystal structures)^30-33^ or low resolution (i.e., examining (sub-)network properties at the protein level, not the residue level)^34-37^. We show utilities of our innovative hotspot-affected interactome network in uncovering novel relationships among different hotspots, generating hypotheses of how hotspot mutations function at the molecular level, and identifying novel oncogenic proteins that do not harbor hotspot mutations themselves. By further complementing the hotspot-affected interactome network with tissue-specific transcriptomes, we prioritize clinically significant hotspots for cancer prognosis and drug development. Together, we offer a biologically-informed framework that characterizes the functional relevance of mutational hotspots and nominate new cancer proteins across human cancers, with interaction-specific resolution at the full proteome scale.

**Fig. 1:**
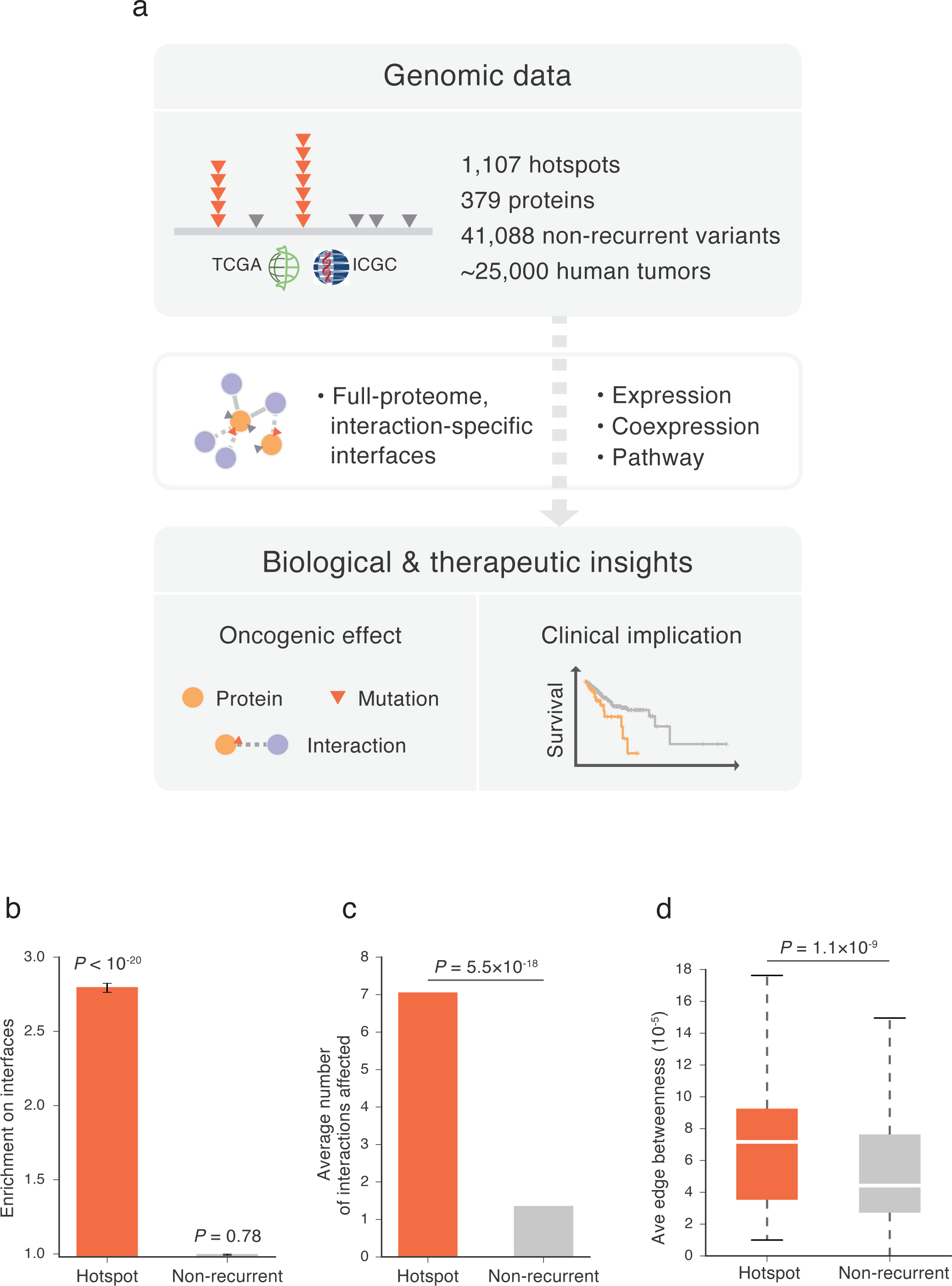
Proteome-wide structural analysis of mutational hotspots on proteins. **a**, Data resource of mutational hotspots and workflow of our full-proteome interaction-specific characterization framework. **b**, Distribution of hotspots and non-recurrent variants on proteins with regard to protein interaction interfaces. Enrichment was calculated as the ratio of the observed fraction of hotspots/variants that occur on interaction interfaces over the fraction of interface residues on corresponding proteins (expected fraction). *P* values were calculated using a two-tailed exact binomial test. The error bars indicate standard error. **c**, Average number of protein interactions affected by hotspots and non-recurrent variants. The number of affected interactions per hotspot/variant was modeled with a negative binomial. **d**, Average edge betweenness of interactions affected by hotspots and non-recurrent variants. Edge betweenness of an interaction was calculated as the sum of the fraction of all-pairs shortest paths that pass through that interaction in the interactome network (Methods). *P* values were calculated using a two-tailed *U*-test.

## Results

### Many hotspot mutations function through affecting specific protein-protein interactions

We previously reported that inherited in-frame disease mutations are enriched on protein interaction interfaces and that alteration of specific protein interactions is critical in the pathogenesis of many disease genes^21^. To explore where somatic hotspots reside with respect to protein interfaces, we mapped 1,107 hotspots (**Supplementary Table 1**) detected from ∼25,000 tumors onto the human interactome (a comprehensive set of 59,073 high-quality physical interactions compiled in HINT^38^ from eight widely used interaction databases^39-46^), in which the interface of each interaction is mapped at residue resolution using Interactome INSIDER^25^. We found that these hotspots are highly enriched on protein interfaces: while interaction interfaces cover 11.0% of the proteins harboring these hotspots, 30.8% of the hotspots fall in interaction interfaces (2.8-fold, *P* = 5.0×10^−62^ by a two-tailed exact binomial test, **Fig. 1b**; Methods). In comparison, when we examined the distribution of ∼40,000 non-recurrent somatic variants (presumed to be predominantly passengers) on the same sets of proteins that harbor hotspots, no enrichment was observed (11.0%, 1.0-fold, *P* = 0.78). This sharp contrast indicates that many hotspot mutations exert their functional effects by affecting specific protein-protein interactions, and that mutation locating on an interaction interface is an important feature that can be used to identify driver mutations.

Leveraging the partner-specific information in our 3D interactome network, we then investigated which and how many interactions are affected by each hotspot. Modeling the count of interactions per hotspot with a negative binomial model yielded a significantly higher number of interactions affected by hotspots than non-recurrent variants on the same set of proteins (means: 7.0 versus 1.3, 5.2-fold, *P* = 5.5×10^−18^, **Fig. 1c**; Methods), suggesting that hotspot mutations preferentially affect “hub interfaces” that involve in a large number of interactions. We further examined the topological positions of these hotspot-affected interactions in the interactome network. Using edge betweenness, where a higher betweenness value indicates more information follow through the corresponding interaction, we found that hotpot-affected interactions on average, have a significantly higher betweenness than those affected by non-recurrent variants (medians: 7.2×10^−5^ versus 4.4×10^−5^, 1.6-fold, *P* = 1.1×10^−9^ by a two-tailed *U*-test, **Fig. 1d**; Methods). Overall, hotspot mutations tend to affect key proteins (high degree) and interactions (high betweenness) that are of great importance topologically for the whole interactome network.

### Hotspot-affected interactions help infer molecular mechanisms of oncogenic hotspots

To assess the oncogenic potential of the hotspots on protein interaction interfaces and the corresponding interactions they affect, we first examined how they link to previously identified cancer genes. Intersecting genes harboring interface hotspots with a list of cancer genes curated in Cancer Gene Census^47^, we found strikingly, ∼80% (79/96) of them are known cancer genes. This gave a 6.8-fold increased odds comparing to that of genes harboring non-interface hotspots (82/202, *P* = 3.7×10^−12^ by a one-tailed Fisher’s exact test, **Fig. 2a** and **Supplementary Table 2**). We next examined the interaction partners and the interaction pairs affected by interface hotspots (see schematics in **Fig. 2b**). There was again significant enrichment for both hotspot-affected interactors and interactions, comparing to the unaffected ones (interactor: 190/1444 versus 150/1795, OR=1.7, *P* = 5.5×10^−6^; interaction: 285/2043 versus 309/3736, OR=1.8, *P* = 1.5×10^−11^; **Fig. 2b** and **Supplementary Table 2**). Overall, these results point to the association of hotspot-affected interaction pairs – involving not only proteins harboring the hotspots but also their affected interaction partners that do not harbor hotspots – with cancer.

**Fig. 2:**
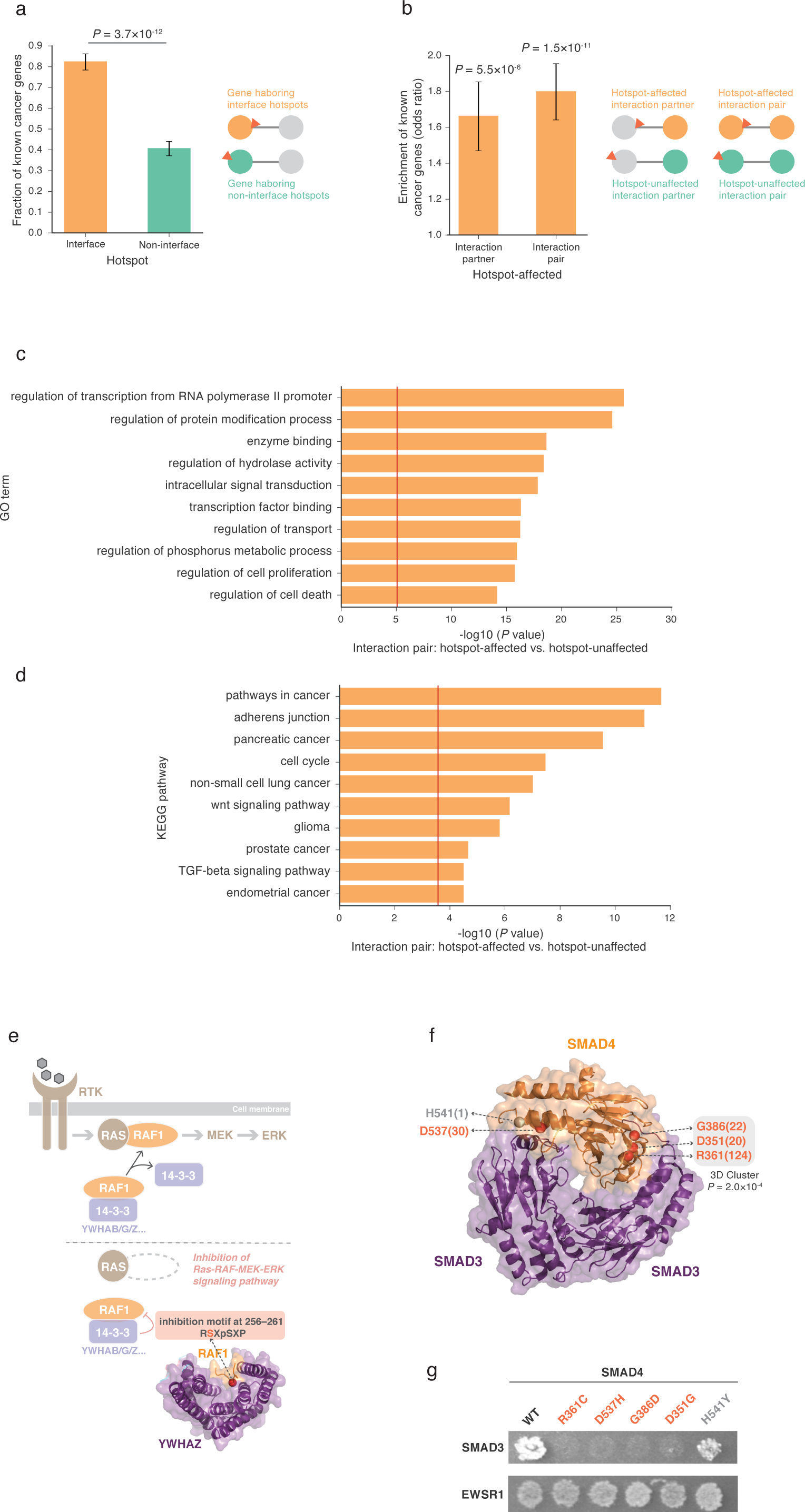
Oncogenic potential of interface hotspots and hotspot-affected interactions. **a**, Association of genes harboring interface and non-interface hotspots with previously known cancer genes. *P* values were calculated using a one-tailed Fisher’s exact test. **b**, Association of hotspot-affected interaction partners and interaction pairs with known cancer genes. *P* values were calculated using a one-tailed Fisher’s exact test comparing the fraction of hotspot-affected interaction partners/pairs that are known cancer genes over that of hotspot-unaffected interaction partners/pairs. An interaction pair was counted when both the gene carrying hotspot and its interaction partner are known cancer genes. **c**,**d**, Gene set enrichment analysis of hotspot-affected interactions. For each (**c**) Gene Ontology (GO) or (**d**) KEGG pathway gene set, enrichment was calculated by comparing the fraction of hotspot-affected interaction pairs that occur in the gene set over that of hotspot-unaffected interaction pairs. An interaction pair was counted when both the gene carrying hotspot and its interaction partner are in the gene set. *P* values were calculated using a one-tailed Fisher’s exact test. The red vertical line indicates statistical significance threshold after Bonferroni correction for 5,917 GO terms and 186 KEGG pathways, respectively. **e**, Implication of hotspot-affected interaction RAF1 S257 – [14-3-3] in oncogenic RAS-RAF-MEK/ERK pathway. A cocrystal structure of RAF1-YWHAZ (PDB ID: 4IHL) highlighting the RAF1 S257 interface hotspot is shown. **f**, Cocrystal structure of SMAD4-SMAD3 trimer (PDB ID: 1U7F) highlighting four SMAD4 interface hotspots (red) and one non-recurrent variant (grey). Number in parentheses indicates the recurrence of corresponding hotspot across ∼25,000 tumor samples. **g**, Effects of SMAD4 hotspot- and non-recurrent mutations on SMAD4 interactions tested by yeast two-hybrid (Y2H) assay. Hotspot mutations (red) disrupted SMAD4-SMAD3 interaction while left SAMD4-EWSR1 interaction intact. The non-recurrent variant (grey) disrupted neither SMAD4-SMAD3 nor SMAD4-EWSR1 interaction.

To further investigate the functional relevance of hotspot-affected interactions in cancer, we performed gene set enrichment analysis asking in what biological processes the pairs of proteins affected by hotspots are functioning together, using unaffected protein pairs as the counterpart (Methods). We found that hotspot-affected interactions are frequently involved in processes that have been tightly linked to cancer (e.g., regulation of transcription, intracellular signal transduction, cell proliferation/death; **Fig. 2c**) and are strongly enriched in curated sets of cancer-associated pathways (**Fig. 2d**). Consequently, alteration of these interactions may be critical for the oncogenicity of hotspot mutations on corresponding interfaces. For example, RAF1 is a serine/threonine kinase with an established role in activating the oncogenic RAS-RAF-MEK/ERK pathway^48^, and its kinase activity can be inhibited by 14-3-3 proteins^49^ (**Fig. 2e**). RAF1 S257L is a hotspot mutation that accounts for 25 and 13% RAF1 substitutions in bowel and lung cancers, respectively. Our framework predicted this hotpot as an interface residue mediating RAF1’s interaction with 14-3-3 proteins. We would then propose RAF1 S257 mutations disrupt RAF1-[14-3-3] interaction, deprive kinase inhibition of RAF1, and thereby potentiate RAS-RAF-MEK/ERK signaling to fuel cancer development. This hypothesis is supported by independent experiments which observed that mutant RAF1^S257L^ lost 14-3-3 binding^50^ and was able to induce anchorage-independent cell growth^51^. Therefore, RAF1 S257 mutations are likely to cause oncogenesis through a “gain of cellular function” via a “loss of molecular inhibition” mechanism. This example helps demonstrate how identification of specific protein interactions affected by a hotpot can generate mechanistic hypothesis about its oncogenicity.

Recurrence of multiple hotspots affecting the same interaction can lend strong evidence to the oncogenic potential of the interaction and shared molecular basis of the hotspots. For instance, we found the most highly recurrent SMAD4 hotspots R361, D537, G386, and D351 all fall in the trimer interface with SMAD3 and three (R361, G386, D351) are significantly clustered in 3D space (*P* = 2.0×10^−4^ using a bootstrapping method by mutation3D^52^, **Fig. 2f**). We experimentally tested the functional impact of these hotspot mutations and the results showed that all mutations disrupted SMAD4-SMAD3 interaction (**Fig. 2g**; Methods). Importantly, these mutations still retained SMAD4’s interaction with EWSR1 (**Fig. 2g**). Thus, the SMAD4 hotspot mutations are likely to function by disrupting specific interaction with SMAD3 rather than causing a loss of the whole protein. We additionally tested a non-recurrent variant on SMAD4-SMAD3 interface (H541Y; **Fig. 2f**) and we found it disrupted neither of the tested interactions (**Fig. 2g**). This suggests that while passenger mutations may still randomly occur on the interfaces, they are much less likely to cause perturbations to protein interactions. Therefore, our framework that identifies interactions affected by mutational hotspots is an effective way to dissect functional consequences and molecular mechanisms of candidate driver mutations in cancer.

### The proteome-scale hotspot-affected interactome network helps identify novel cancer proteins without hotspot mutations

Given the functional significance and oncogenic potential of hotspot-affected interactions, we constructed a “hotspot-affected interactome network” by connecting all 2,083 interactions affected by hotspots at the whole proteome scale (**Fig. 3a** and **Supplementary Table 2**). As a control, the remaining 3,736 unaffected interactions of proteins harboring hotspots were built into another network, referred to as the hotspot-unaffected network (**Fig. 3a**). We first examined how known cancer-associated proteins distribute between and within these two networks. Overall there was a 1.2-fold enrichment for cancer proteins in the hotspot-affected network versus the unaffected network (210/1471 versus 307/2617, *P* = 0.01 by a one-tailed *Z*-test). Interestingly, with an increasing network degree (number of interactions associated with a protein), the enrichment became increasingly stronger (**Fig. 3b** and **Supplementary Table 3**), suggesting that cancer proteins tend to act as hubs in the hotspot-affected network.

**Fig. 3:**
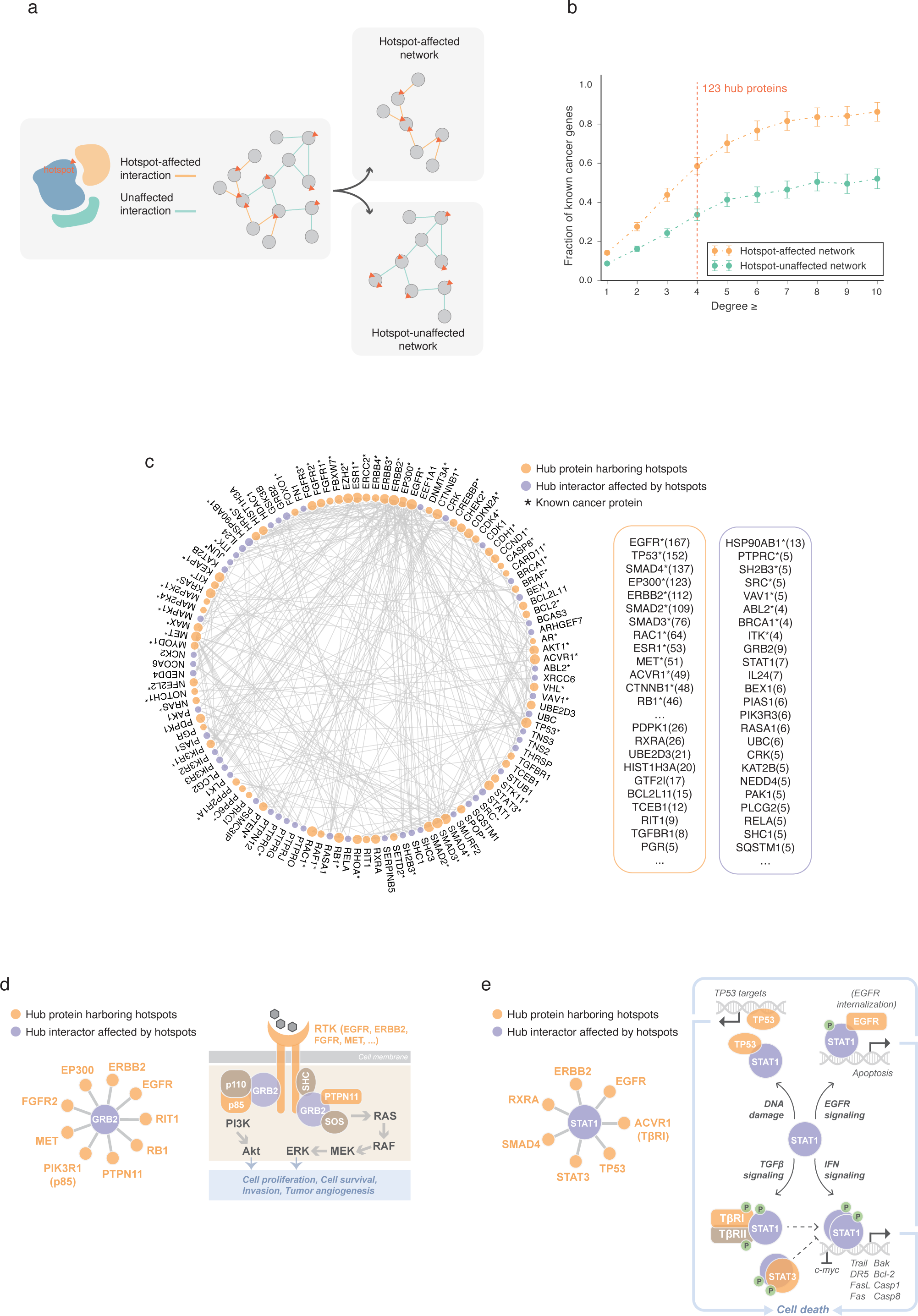
Identification of novel cancer proteins using our hotspot-affected network. **a**, Schematic illustration of constructing hotspot-affected and hotspot-unaffected networks. **b**, Association of proteins in the hotspot-affected and hotspot-unaffected networks with previously known cancer proteins. The error bars indicate standard error. **c**, A network view of hub proteins prioritized by our hotspot-affected network. Proteins that harbor hotspots are shown in orange and proteins with no hotspots are shown in purple. Listed proteins are ranked first by whether the protein is a known cancer protein (indicated by asterisk) and then by their degree in the hotspot-affected network (shown in parentheses). See supplementary table 3 for a full list. **d**,**e**, Implication of the top-ranked hub proteins (**d**) GRB2 and (**e**) STAT1 in oncogenesis.

We prioritized 123 hubs (degree≥4) from the hotspot-affected network as candidate cancer proteins (**Fig. 3c** and **Supplementary Table 3**). As expected, topping the list are well-known cancer proteins such as epidermal growth factor receptors (EGFR, ERBB2), tumor protein p53, and SMAD family of signal transduction proteins. An important feature of our network-based approach, rather than single-protein level functional predictions, is its ability to discover proteins that do not harbor any hotspots themselves, but are frequently targeted by hotspot-affected interactions. While these proteins may go unrecognized by other methodologies due to the lack of recurrent mutations, we interpret them as crucial nodes that connect many different hotspots and may serve as a convergence point for their oncogenic effects and thus are candidate therapy targets. Among such hubs, GRB2 ranked as the top novel candidate which links to nine proteins harboring hotspots (“hotspot-interactors”; **Fig. 3d**). GRB2 acts as an adapter protein between cell surface receptors and intracellular signal transducers. Correspondingly, we found four GRB2 hotspot-interactors are receptor tyrosine kinases (RTKs: EGFR, ERBB2, FGFR2, MET) and two are downstream effectors (PTPN11, p85) in pathways largely involved in human cancers (**Fig. 3d**). Consequently, GRB2 appears as the pivot for these proteins harboring hotspots to launch oncogenic signaling cascades. Although no genetic alteration on GRB2 has been identified oncogenic, our network-based approach implicates GRB2 as a cancer-associated protein that empowers the oncogenicity of many hotspot mutations on its interacting proteins. Meanwhile, the hub role of GRB2 makes it an attractive target for cancer therapeutics; antagonists of GRB2 activity^53^ can potentially benefit groups of patients who carry mutations affecting interactions with GRB2.

Besides acting as a convergence point, a hub could alternatively mediate diverse oncogenic activities when affected by different hotspots. Signal transducer and activator of transcription (STAT) 1, the second ranked candidate in our network, represents one such instance. As a signal transducer, STAT1 can mediate TGF-β signaling by directly interacting with TGF-β receptors^54^(**Fig. 3e**). As an activator of transcription, STAT1 regulates a number of genes, either in a form of homo/heterodimer best known in the response to interferons (IFNs)^55^, or by cooperating with other proteins such as TP53 and EGFR in p53/EGF-induced apoptotic pathways^56-59^ (**Fig. 3e**). Interestingly, while serving as components of different pathways, these interactions all point to STAT1’s function in promoting cell death. For instance, STAT1 works cooperatively with tumor suppressor p53 to induce apoptosis^56-58^ (**Fig. 3e**) while performs oppositely with STAT3 – a known oncoprotein – where STAT1 up/downregulating pro/anti-apoptotic genes is suppressed by STAT3^55^ (**Fig. 3e**). Therefore, our results support a tumor suppressor role for STAT1 and we implicate specific hotspot-affected interactions that may underlie its tumor-suppressive function. Moreover, we would predict genetic alterations that cause a loss or reduction of STAT1 to have oncogenic effects. While no STAT1 truncating mutations have been tested directly, STAT1 knockout mice have been shown to spontaneously develop mammary carcinomas^60^. Together, we expect that as more mutations are uncovered and studied, more nodes and edges will be added to our hotspot-affected network, providing complementary evidence that strengthens previously identified associations and enhances the discovery of new ones.

### Cancer-type-specific hotspots affect different interactions in different cancers

In Mendelian disorders, it has been shown that mutations on the same protein can cause distinct diseases through altering different protein interactions^21,23^. To explore the association of hotspots and hotspot-affected interactions with different cancers, we analyzed the subset of hotspots detected from individual cancer types^14^. In total, 719 unique cancer type-specific hotspots were identified, of which ∼30% occurred in more than one cancer type (“multi-cancer hotspots”; **Fig. 4a** and **Supplementary Table 4**). We first examined how these hotspots distributed in the interactome network and found that multi-cancer hotspots, comparing to single-cancer hotspots, tend to affect more central positions (degree medians: 35 versus 22.5, fold-change (FC) = 1.6, *P* = 5.8×10^−9^ by a one-tailed *U*-test, **Fig. 4b**). We next determined multi-cancer interactions, as those affected by either one or more multi-cancer hotspots or multiple single-cancer hotspots of different cancers (**Supplementary Table 4**). In line with the centrality of multi-cancer hotspots, we found multi-cancer interactions also tend to occupy central positions within the interactome network, as indicated by their high edge betweenness compared to that of single-cancer interactions (medians: 7.5×10^−^ 5 versus 5.9×10^−5^, FC = 1.3, *P* = 3.1×10^−7^, **Fig. 4c**). These findings suggest that the multi-cancer roles of hotspots and hotspot-affected interactions may arise from their network centrality, which allows them to function through a wide range of downstream targets.

**Fig. 4:**
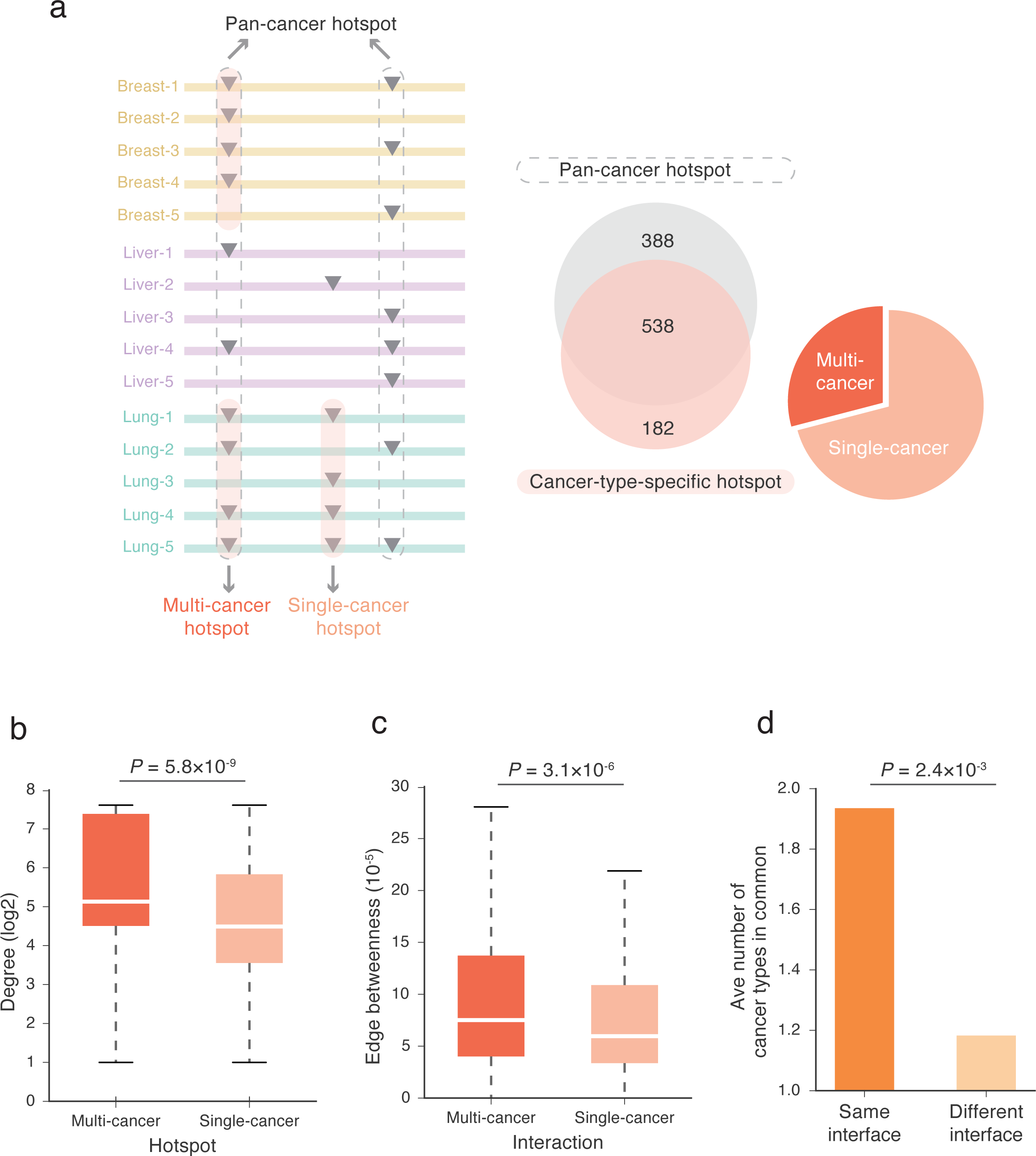
Association of cancer type-specific hotspots with specific protein interactions. **a**, Categorization of pan-cancer versus cancer type-specific hotspots and multi-cancer versus single-cancer hotspots. **b**, Degree distributions of proteins harboring multi-cancer and single-cancer hotspots. Degree values are transformed by log2 for presentation purposes. *P* values were calculated using a one-tailed *U*-test. **c**, Edge betweenness distributions of multi-cancer and single-cancer interactions. *P* values were calculated using a one-tailed *U*-test. **d**, Average number of cancer types shared between hotspots on the same interface and between hotspots on different interfaces. The number of shared cancer types between each pair of hotspots was modeled with a negative binomial.

We then asked whether hotspots targeting different interaction partners tend to be involved in different cancers. By comparing the number of cancer types shared by hotspots on the same protein, we found a significantly greater number of common cancer types shared by hotspots affecting the same interactions than those affecting different interactions (1.6-fold, *P* = 3.7×10^−4^ by a negative binomial model, **Fig. 4d**; Methods). The result indicates that hotspots affecting different interactions, while on the same protein, are likely to involve in different cancers. This reinforces our notion that alteration of specific protein interactions is critical for the oncogenicity of hotspot mutations. Therefore, linking specific interactions to hotspots of specific cancer types – that is, constructing a cancer-type-specific, hotspot-affected network – would further delineate functional hotspots and interactions and better understand their molecular mechanisms in specific cancers.

### Cancer-type-specific hotspot-affected interactome networks prioritize hotspots for prognosis and drug development

Many interactions may not happen across all tissues, and consequently, for a particular type of cancer, a hotspot-affected interaction is only meaningful if that interaction happens in the corresponding tissue. Since tissue specificity is largely driven by the epigenetic landscape across tissues^61^, we surveyed the expression patterns of genes encoding hotspot-affected interacting proteins, using RNA-seq data from matched tumor-adjacent normal tissues (Methods). We observed that genes encoding hotspot-affected interaction pairs, on average, have a significantly higher expression correlation than genes encoding hotspot-unaffected pairs (medians: 0.27 versus 0.23, FC = 1.2, *P* = 1.6×10^−6^ by a one-tailed *U*-test, **Fig. 5a**). This agrees well with our expectation that the pairs of proteins affected by hotspots are functionally associated with each other. We then incorporated the tissue-specific gene coexpression to construct cancer-type-specific, hotspot-affected interactome networks for all cancer types where data are available. Specifically, for each cancer type, we only include hotspot-affected interactions where both interacting partners are co-expressed in the corresponding tissue (**Fig. 5b**; Methods).

**Fig. 5:**
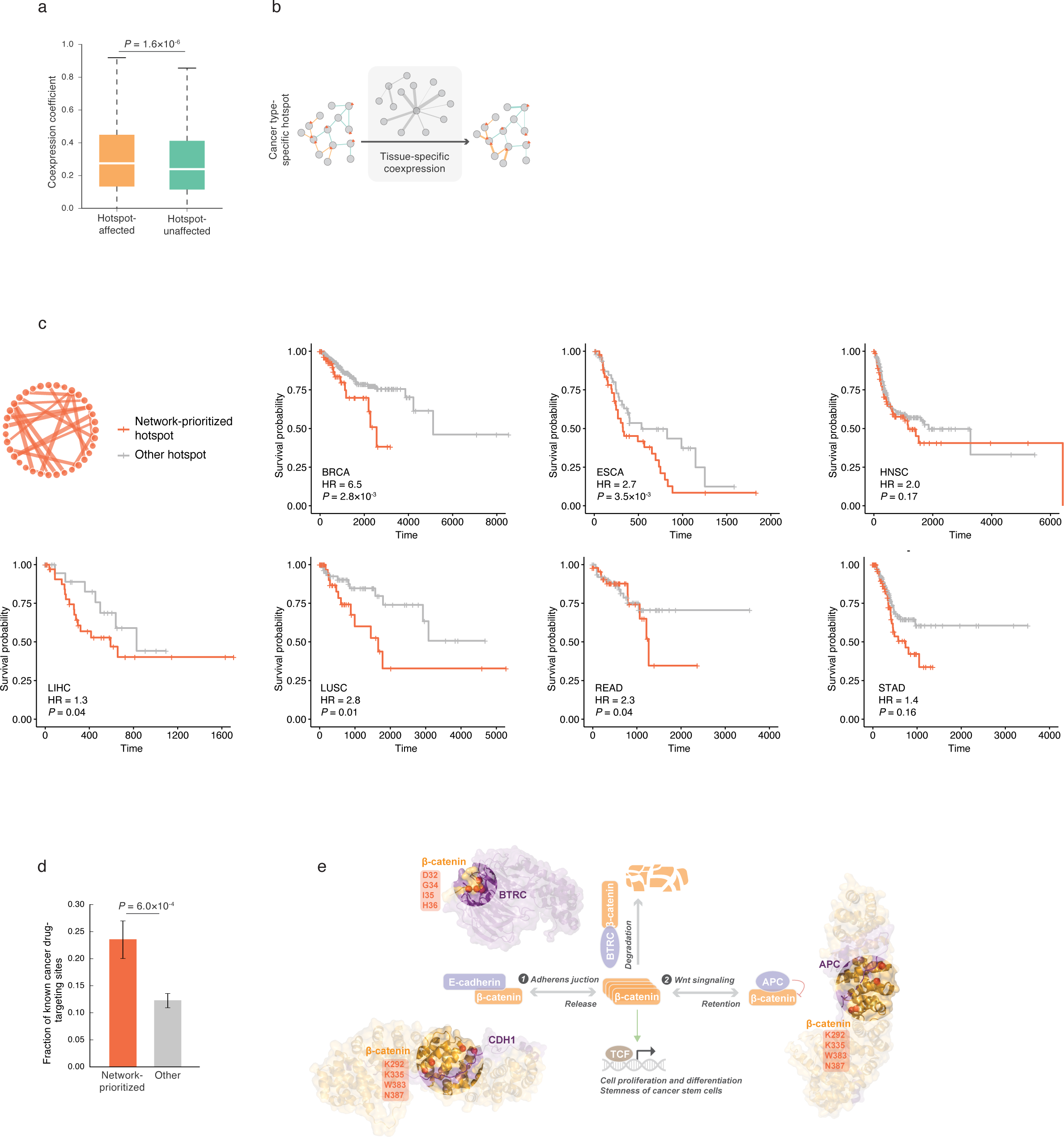
Construction of cancer type-specific hotspot-affected networks and their clinical utilities. **a**, Tissue-specific coexpression levels of genes encoding hotspot-affected and -unaffected interaction pairs in corresponding cancer types. *P* values were calculated using a one-tailed *U*-test. **b**, Schematic illustration of constructing a cancer type-specific hotspot-affected network. For a particular type of cancer, cancer type-specific hotspot-affected interactions were first determined, and for each interaction pair, gene coexpression coefficient was calculated from matched tumor-adjacent normal tissue; pairs with an absolute Pearson correlation coefficient >0.5 were selected for constructing the ultimate network. **c**, Association of our network-prioritized hotspots with patients’ survival. Survival probabilities were estimated using Kaplan-Meier method for patients carrying network-prioritized hotspots and patients carrying other hotspots. Hazard ratio (HR) and *P* values were calculated using a Cox regression model using network-prioritized hotspot as the predictor and controlling for clinical covariates including patient age, gender, tumor stage, and subtype. BRCA: breast invasive carcinoma (*n* = 60 for network-prioritized, *n* = 398 for other); ESCA: esophageal carcinoma (*n* = 44, 52); HNSC: head and neck squamous cell carcinoma (*n* = 76, 186); LIHC: liver hepatocellular carcinoma (*n* = 38, 25); LUSC: lung squamous cell carcinoma (*n* = 42, 64); READ: rectum adenocarcinoma (*n* = 47, 67); STAD: stomach adenocarcinoma (*n* = 88, 124). **d**, Association of our network-prioritized hotspots with known cancer drug targets. *P* values were calculated using a one-tailed *U*-test. **e**, Implication of β-catenin as a drug target for patients carrying β-catenin hotspots. Cocrystal structures of β-catenin – APC (PDB ID: 1TH1) and β-catenin – BTRC (PDB ID: 1P22), and a homology model of β-catenin – CDH1 (template PDB ID: 1I7W), highlighting β-catenin interface hotspots are shown.

To evaluate the utility of our cancer-type-specific hotspot-affected interactome networks, we investigated whether the network-prioritized hotspots are associated with patients’ clinical outcomes. In each of the seven cancer types that had sufficient network and expression data, we compared the progression-free survival between patients carrying hotspot mutations, grouped by whether the hotspot is prioritized by our network or not (**Supplementary Table 5**). Kaplan-Meier estimates revealed shorter survival time in patients carrying network-prioritized hotspot mutations than patients carrying other hotspot mutations, across all cancer types (**Fig. 5c**). We then employed Cox regression to assess the prognostic potential of network-prioritized hotspot, controlling for clinical covariates including patient age, gender, tumor stage, and subtype (Methods). The model yielded significant associations between network-prioritized hotspots and poor survival outcomes for patients (hazard ratio (HR) > 1.0 and *P* < 0.05, in 5/7 cancer types; **Fig. 5c**). Thus, hotspots prioritized by our networks could serve as prognostic biomarkers that can be used for progression prediction and therapy selection for induvial patients.

To further evaluate the utility of our network-prioritized hotspots in targeted therapy, we surveyed a list of 287 genetic biomarkers that are targeted by known cancer drugs (curated in My Cancer Genome: mycancergenome.org). We found a significant enrichment of our prioritized hotspots as known drug targets compared to other hotspots (35/149 versus 76/622, OR = 2.2, *P* = 6.0×10^−4^ by a one-tailed *U*-test, **Fig. 5d**). This reinforces the functional significance of our prioritized hotspots in cancer progression, and thereby these hotspots may provide a valuable list of “yet-to-be-drugged” candidates for new drug development. Among them, β-catenin emerged attractive as it harbors a number of prioritized hotspots (D32, G34, I35, H36, K292, K335, W383, N387). Interestingly, these hotspots were predicted to affect two distinct interfaces involved in different pathways yet they appeared to function interactively in controlling the level of β-catenin (**Fig. 5e**). Therefore, targeting β-catenin could be a promising therapeutic strategy for patients with mutations on these hotspots. In fact, although currently no β-catenin-targeted therapies are being used for treatment, β-catenin status serves as an inclusion eligibility criteria in several clinical trials (NCT02013154, NCT02724579, NCT02066220). These findings suggest that our prioritization would have valid and immediate clinical implications. Collectively, we demonstrate that our framework of focusing on hotspot-affected interactions and networks provides an effective way to prioritize for biological and clinical studies of functional and potentially druggable hotspots.

Since high recurrence has been regarded as a hallmark of cancer-driving mutations, it is natural for researchers and clinicians to prioritize hotspots by their degree of recurrence. To examine the value of our network-based prioritization in providing new independent information, we divided the hotspots into two groups by their statistical ranking (upper 50% and lower 50%) and repeated all analyses separately within each group. Intriguingly, we found all results remained the same for both upper- and lower-ranked hotspots (**Supplementary Fig. 1a-1i**). This clearly shows that our network-based prioritization can provide orthogonal information in addition to recurrence to better guide functional studies and clinical interpretations. In fact, recurrence by itself is not informative enough to distinguish clinically relevant hotspots (**Supplementary Fig. 2**). Collectively, we suggest that one key feature to inform the functional relevance of cancer hotspots is whether they fall in protein interfaces that mediate important interactions. We present a scalable and reliable framework that identifies specific protein-protein interactions affected by hotspots at the full-proteome scale and yields mechanistic and clinical insights towards how a hotspot mutation may contribute to tumorigenesis.

## Discussion

Cancer driver mutations are frequently nominated based on their recurrence rate in different cancers. However, because there are many additional factors that affect mutation rates^16,62^, not all hotspot mutations are cancer drivers. Here we demonstrated that many cancer hotspot mutations function through affecting specific protein-protein interactions and that our full-proteome, interaction-specific 3D interactome network framework can effectively identify functional hotspots and hotspot-affected interactions. Because experimental examination of hotspots is limited in scale and current bioinformatics tools hardly interpret oncogenicity, our framework makes significant contributions in systematically prioritizing and mechanistically characterizing hotspots in human cancers. Our results revealed that hotspots and hotspot-affected interactions preferentially occupy central positions in the interactome network and frequently target previously known cancer proteins and pathways. While we suggested that identifying specific individual interactions affected by a hotspot can already help understand its oncogenicity, additional functional associations were uncovered when we connected individual hotspot-affected interactions together into a network. This hotspot-affected network led to the discovery of novel cancer proteins which themselves do not harbor hotspots but serve as important nodes for hotspots on their interaction partners to function. Therefore, although our analyses started with ∼1,000 published hotspots, our framework expanded the scope of existing datasets and can be readily applied as more cancer mutations are identified. Finally, we showed clinical utilities of applying our framework in specific cancer types towards improving prognosis and therapy targets.

One unique strength of our framework resides in its full-proteome scale prediction of interfaces for interactions with no cocrystal structures or homology models available. While we emphasize the scalability of our framework, we ensure the reliability of our findings by repeating all analyses using only protein interfaces resolved from cocrystal structures and homology models. All results agreed well with those calculated from full interface data yet had reduced statistical significance due to limited sample size (**Supplementary Fig. 3a-3j**). These results not only confirm the validity of our findings in this study, but also underscore the effectiveness of our full-proteome framework in characterizing mutational hotspots. Moreover, we recognize that the current human interactome may be subject to sampling bias from small-scale studies^38,63^, where certain proteins (e.g., TP53) may have been studied intensively against a great number of interactors. To address this issue, we re-examined the network properties of hotspots and hotspot-affected interactions using only high-throughput-derived human interactions^38^, and our results on network centralities remain unchanged (**Supplementary Fig. 4a-4c**), further confirming the importance of using network properties to interpret the functional significance of hotspots in cancer.

Taken together, our innovative network-based framework provides an effective way to identify candidate driver hotspots, and to dissect the molecular mechanisms underlying their oncogenicity. Our findings would help researchers and clinicians nominate hotspots that can serve as biomarkers for cancer prognosis in individual patients, personalized treatment, and development of new therapeutics.

## Methods

### Enrichment of hotspots and non-recurrent variants on protein interfaces

For each hotspot or non-recurrent variant, with each of its interaction partners, we considered it to be on the interface for this specific interaction if it has a probability score of high or very high in Interactome INSIDER prediction. Suppose the locations of hotspots are not influenced by the interface architecture of the protein, then their relative length should determine the frequency of hotspots on interfaces. The fraction of hotspots expected by chance on interfaces was calculated by adding the total sequence length of interface residues in all proteins harboring hotspots, and dividing it by the length of all proteins combined; let the probability of falling in an interaction interface be *p*. Let the number of observed hotspots falling in the interfaces be *S*, and let *N* be the total number of hotspots. An exact binomial test was then computed from *p, S* and *N*. CIs were based on the 95% CI for an exact binomial, and then transformed to the risk ratio (enrichment) using the expectation in the denominator and the lower/upper bound in the numerator.

The set of 251 proteins harboring hotspots and containing at least one interface residue was included in the interface enrichment calculations. The total length of 251 proteins is 202,578 and the total number of interface residues on these proteins is 22,372, then the probability for a hotspot or non-recurrent variant to fall in interfaces was computed to be 11.0%. Of the 966 hotspots observed on these proteins 298 fell in interfaces (30.8%), yielding a significant enrichment of 2.8 fold (2.5-3.1 95% CI, *P* = 5.0 × 10^−62^). In contrast, the 28,657 non-recurrent variants occurred on interfaces of these proteins with the expected rate (3,149/28,657 = 11.0%, enrichment = 1.0 [0.96-1.03 95% CI], *P* = 0.78).

### Modeling the number of interactions as a function of hotspot status

Some hotspots can affect *M* = 1, 2, …, *I* interactions while others do not affect any interactions, *M* = 0. To account for the dispersion in *M*, and to determine whether *M*is stochastically greater for hotspot than non-recurrent variant, we modeled *M* as a negative binomial distribution and fit it to hotspot status (“1” for hotspot and “0” for non-recurrent variant).

### Computing edge betweenness

We computed edge betweenness for protein interaction pairs (edges) in the interactome network using algorithm from Ulrik Brandes^64,65^ (built in Python module NetworkX): betweenness of an edge *e* is the sum of the fraction of all-pairs shortest paths that pass through *e*:

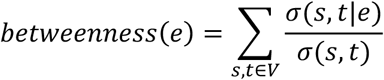

where *V* is the set of nodes, *σ*(*s, t*) is the number of shortest (*s, t*)-paths, and *σ*(*s, t*|*e*) is the number of those paths passing through edge *e*.

### Gene set enrichment analysis

Enrichment of 2,043 hotspot-affected interaction pairs over 3,736 hotspot-unaffected interaction pairs was tested for 15,917 Gene Ontology (GO) and 186 KEGG pathway gene sets (obtained from MSigDB^66^). An interaction pair was counted when both the gene carrying hotspot and its interaction partner are in the gene set under examination. Statistical significance threshold was corrected by Bonferroni.

### Experimental examination of SMAD4 hotspot mutations using yeast two-hybrid (Y2H) assay

To perform Y2H, pDEST-AD and pDEST-DB plasmid vectors corresponding to the GAL4-activating domain (AD) and DNA-binding (DB) domain, respectively, were used. Full-length Clone-seq-identified mutant clones were transferred into Y2H-amenable pDEST-DB and pDEST-AD vectors by Gateway LR reactions and then transformed into *MATα* Y8930 and *MATα* Y8800, respectively. All DB-ORF *MATα* transformants, including wild-type ORFs, were then mated against corresponding wild-type and mutant AD-ORF *MATα* transformants in a pairwise orientation on YEPD agar plates. After mating, yeast was replica-plated onto selective SC–Leu–Trp–His+1 mM of 3-amino-1,2,4-triazole (3AT) as well as SC–Leu–Trp–Adenine plates. Interactions were scored after three days of incubation and five days of incubation for SC–Leu–Trp+3AT and SC–Leu–Trp–Ade plates, respectively. To screen out autoactivating DB-ORFs, all DB-ORF *MATα* transformants were also mated pairwise against empty pDEST-AD *MATα* transformants and scored for growth on SC–Leu–Trp+3AT and SC–Leu–Trp–Ade plates. DB-ORFs that trigger reporter activity under this setup were removed from further experiments.

### Comparing the number of common cancer types shared between hotspots

To test whether hotspots affecting same/different interactions tend to associate with same/different cancer types, we performed pairwise comparison between cancer types linked to hotspots on the same protein. We examined on 26 proteins, 97 pairs of hotspots on the same protein interaction interfaces (i.e., affecting the same set of interactions) and 198 pairs of hotspots on different interfaces (i.e., affecting non-overlapping sets of interactions). The number of cancer types shared by a pair of hotspots was modeled by a negative binomial distribution, using whether the pair of hotspots are on the same interface as predictor (“1” for same interface and “0” for different interface).

### Tissue-specific coexpression analysis

RNA-seq data of tumor-adjacent normal tissues were obtained from Genomic Data Commons (GDC) Data Portal. Ten cancer types (bowel, breast, esophagus, head and neck, kidney, liver, lung, prostate, stomach, and thyroid) with at least 40 normal tissue samples were considered for coexpression analysis. For each cancer type, we calculated coexpression coefficients of genes encoding hotspot-affected and -unaffected interaction pairs across corresponding normal tissue samples. Excluding homodimers, we compared the average absolute Pearson correlation coefficients of 2,414 hotspot-affected and 3,970 hotspot-unaffected interaction pairs. Tissue-specific coexpression networks were constructed using gene pairs with an absolute Pearson correlation coefficient >0.5.

### Survival analysis

We compared the progression-free survival between patients carrying our network-prioritized hotspots and patients carrying other hotspots. Survival curves were generated using Kaplan-Meier estimation. Statistical association between network-prioritized hotspot and patient survival outcome was evaluated using a Cox regression model, controlling for clinical covariates including patient age, gender, tumor stage, and subtype. Patient clinical data was obtained from TCGA Pan-Cancer Clinical Data Resource (CDR)^67^, and cancer types that have at least 50 patients with available clinical information were considered for survival analysis.

## Acknowledgements

We thank the members of the Yu Lab for constructive discussions. This work was supported by National Institute of General Medical Sciences grants (R01 GM104424, R01 GM124559, R01 GM125639); a National Cancer Institute grant (R01 CA167824); a Eunice Kennedy Shriver National Institute of Child Health and Human Development grant (R01 HD082568); a National Human Genome Research Institute grant (UM1 HG009393); a National Science Foundation grant (DBI-1661380); and a Simons Foundation Autism Research Initiative grant (SF367561) to H.Y.

## Authors’ contributions

S.C., X.W., and H.Y. conceived the study. H.Y. oversaw all aspects of the study. S.C. designed and performed all computational analyses with input from Y.Z. and H.Y. Y.L. performed laboratory experiments. S.C. wrote the manuscript with input from S.D.W., S.M.L., and H.Y. All authors edited and approved of the final manuscript.

## Competing interests

The authors declare no competing interests.

