## Supplementary Figures for "A full-proteome, interaction-specific characterization of mutational hotspots across human cancers"

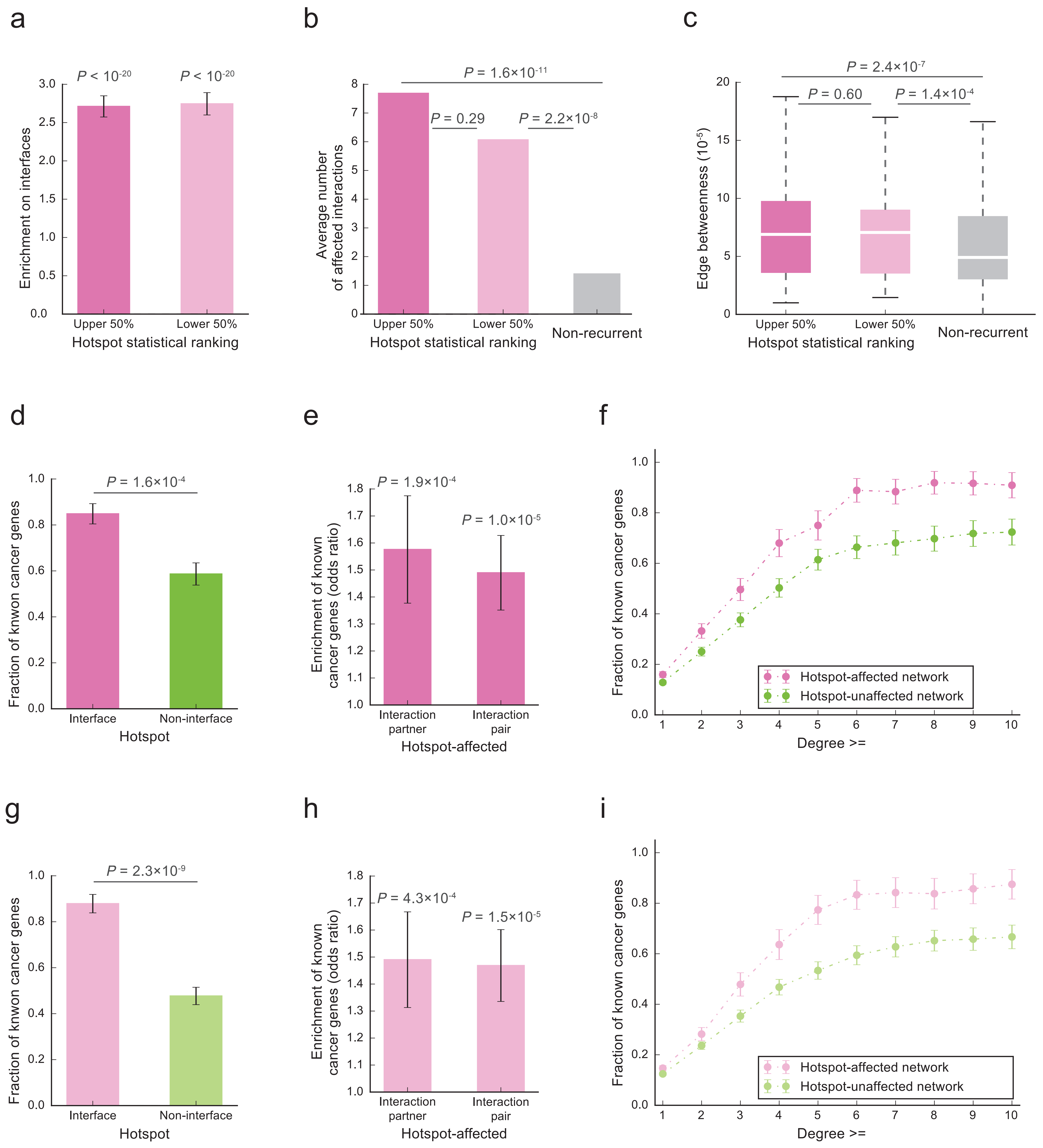


**Supplementary Fig. 1**: Analyses of mutational hotspots grouped by their statistical ranking (dark pink: upper 50%; light pink: lower 50%). **a**, Distribution of upper- and lower-ranked hotspots on proteins with regard to protein interaction interfaces. Enrichment was calculated as the ratio of the observed fraction of hotspots/variants that occur on interaction interfaces over the fraction of interface residues on corresponding proteins (expected fraction). P values were calculated using a two-tailed exact binomial test. The error bars indicate standard error. **b**, Average number of protein interactions affected by upper-/lower-ranked hotspots and non-recurrent variants. The number of affected interactions per hotspot/variant was modeled with a negative binomial. **c**, Average edge betweenness of interactions affected by upper-/lower-ranked hotspots and non-recurrent variants. Edge betweenness of an interaction was calculated as the sum of the fraction of all-pairs shortest paths that pass through that interaction in the interactome network (Methods). P values were calculated using a two-tailed U-test. **d**,**g**, Association of genes harboring interface and non-interface hotspots with previously known cancer genes (**d**/**g**: upper-/lower-ranked hotspots). P values were calculated using a one-tailed Fisher's exact test. **e**,**h**, Association of hotspot-affected interaction partners and interaction pairs with known cancer genes (**e**/**h**: upper-/lower-ranked hotspots). P values were calculated using a one-tailed Fisher's exact test comparing the fraction of hotspot-affected interaction partners/pairs that are known cancer genes over that of hotspot-unaffected interaction partners/pairs. An interaction pair was counted when both the gene carrying hotspot and its interaction partner are known cancer genes. **f**,**i**, Association of proteins in the hotspot-affected and hotspot-unaffected networks with previously known cancer proteins (**f**/**i**: upper-/lower-ranked hotspots). The error bars indicate standard error.

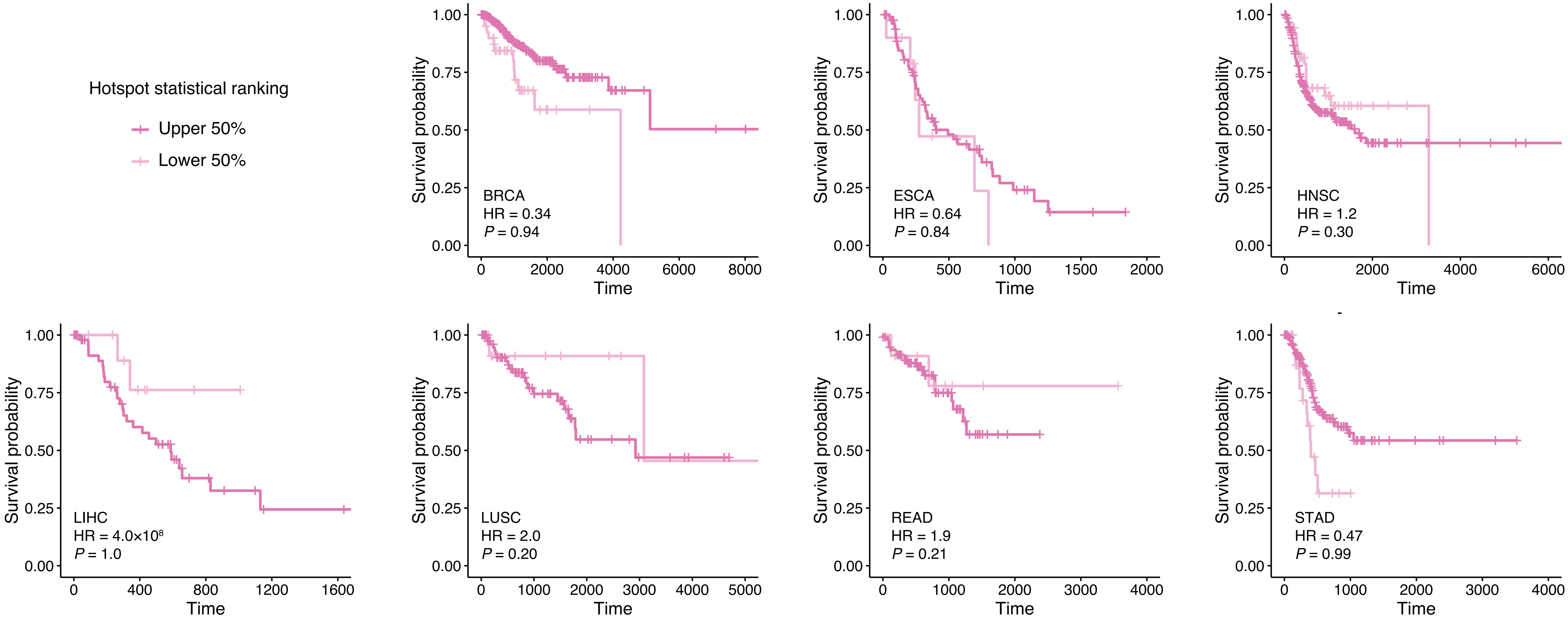


**Supplementary Fig. 2**: Association of the statistical ranking of hotspots with patients' survival. Survival probabilities were estimated using Kaplan-Meier method for patients carrying upper-ranked hotspots and patients carrying lower-ranked hotspots. Hazard ratio (HR) and P values were calculated using a Cox regression model using upper-ranked hotspot as the predictor and controlling for clinical covariates including patient age, gender, tumor stage, and subtype. BRCA: breast invasive carcinoma (n = 414 for upper-ranked, n = 44 for lower-ranked); ESCA: esophageal carcinoma (n = 86, 10); HNSC: head and neck squamous cell carcinoma (n = 210, 52); LIHC: liver hepatocellular carcinoma (n = 52, 11); LUSC: lung squamous cell carcinoma (n = 93, 13); READ: rectum adenocarcinoma (n = 103, 11); STAD: stomach adenocarcinoma (n = 181, 31).

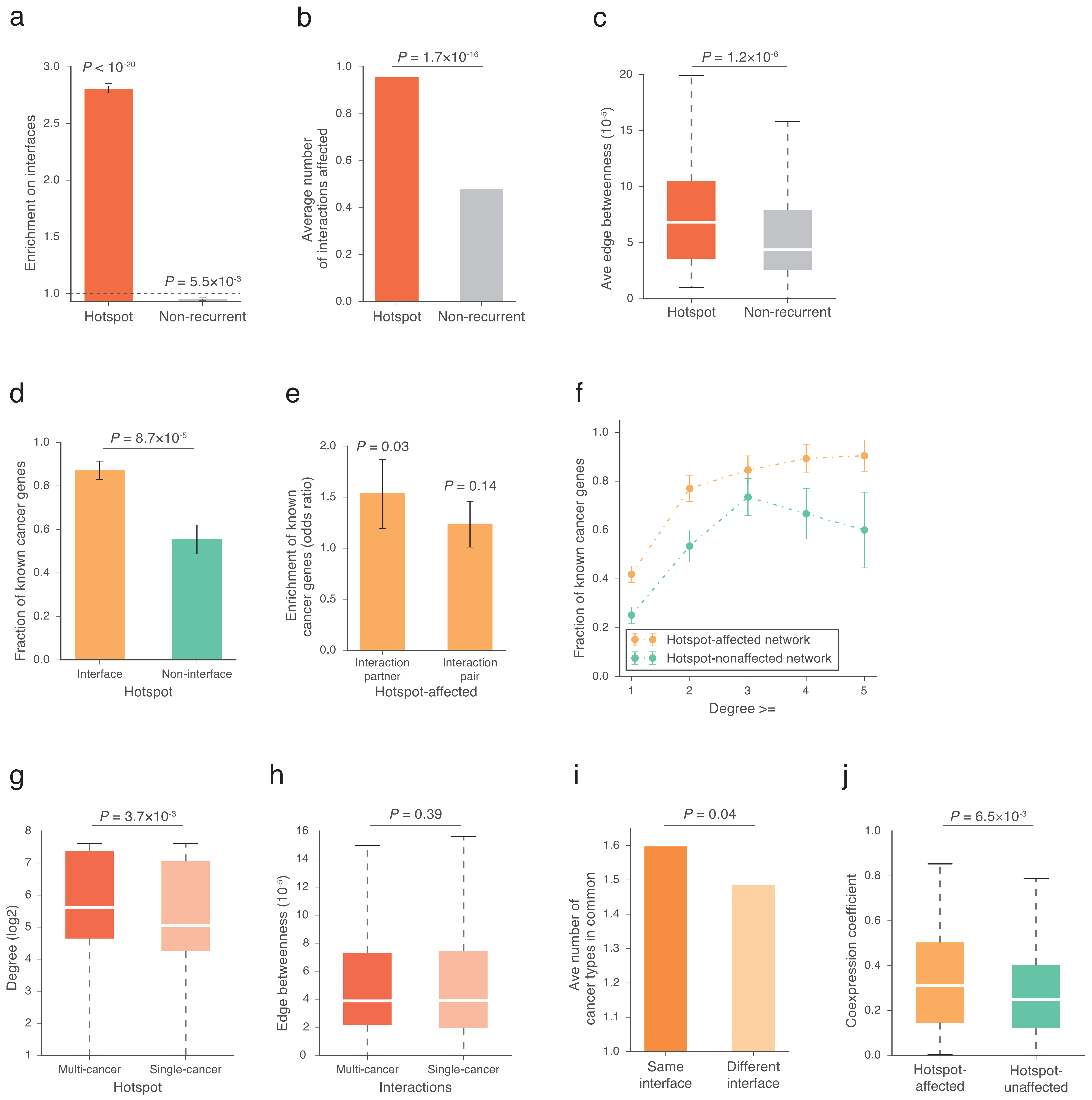


**Supplementary Fig. 3**: Analyses of mutational hotspots using protein interfaces resolved from cocrystal structures and homology models. **a**, Distribution of hotspots and non-recurrent variants on proteins with regard to protein interaction interfaces. Enrichment was calculated as the ratio of the observed fraction of hotspots/variants that occur on interaction interfaces over the fraction of interface residues on corresponding proteins (expected fraction). P values were calculated using a two-tailed exact binomial test. The error bars indicate standard error. **b**, Average number of protein interactions affected by hotspots and non-recurrent variants. The number of affected interactions per hotspot/variant was modeled with a negative binomial. **c**, Average edge betweenness of interactions affected by hotspots and non-recurrent variants. Edge betweenness of an interaction was calculated as the sum of the fraction of all-pairs shortest paths that pass through that interaction in the interactome network (Methods). P values were calculated using a two-tailed U-test. **d**, Association of genes harboring interface and non-interface hotspots with previously known cancer genes. P values were calculated using a one-tailed Fisher's exact test. **e**, Association of hotspot-affected interaction partners and interaction pairs with known cancer genes. P values were calculated using a one-tailed Fisher's exact test comparing the fraction of hotspot-affected interaction partners/pairs that are known cancer genes over that of hotspot-unaffected interaction partners/pairs. An interaction pair was counted when both the gene carrying hotspot and its interaction partner are known cancer genes. **f**, Association of proteins in the hotspot-affected and hotspot-unaffected networks with previously known cancer proteins. The error bars indicate standard error. **g**, Degree distributions of proteins harboring multi-cancer and single-cancer hotspots. Degree values are transformed by log2 for presentation purposes. P values were calculated using a one-tailed U-test. **h**, Edge betweenness distributions of multi-cancer and single-cancer interactions. P values were calculated using a one-tailed U-test. **i**, Average number of cancer types shared between hotspots on the same interface and between hotspots on different interfaces. The number of shared cancer types between each pair of hotspots was modeled with a negative binomial. **j**, Tissue-specific coexpression levels of genes encoding hotspot-affected and -unaffected interaction pairs in corresponding cancer types. P values were calculated using a one-tailed U-test.

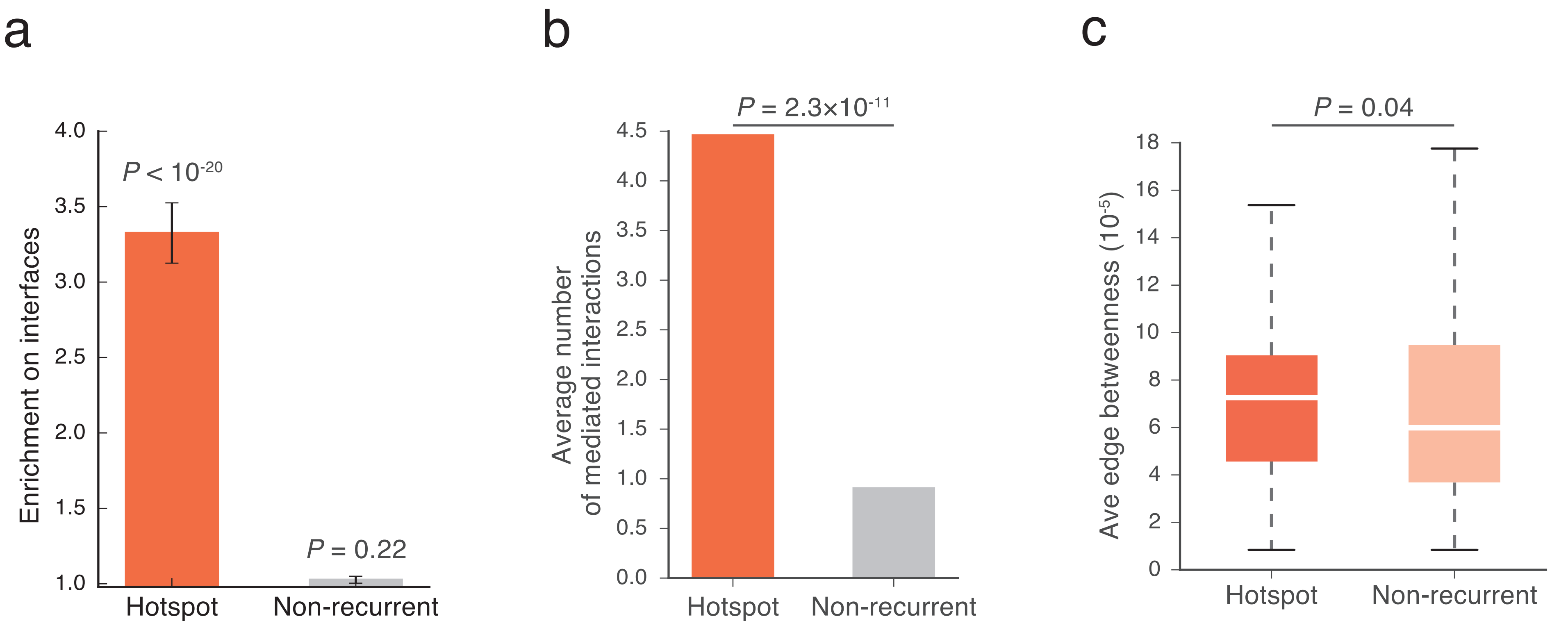


**Supplementary Fig. 4**: Network properties of mutational hotspots in a high-throughput-derived human interactome. **a**, Distribution of hotspots and non-recurrent variants on proteins with regard to protein interaction interfaces. Enrichment was calculated as the ratio of the observed fraction of hotspots/variants that occur on interaction interfaces over the fraction of interface residues on corresponding proteins (expected fraction). P values were calculated using a two-tailed exact binomial test. The error bars indicate standard error. **b**, Average number of protein interactions affected by hotspots and non-recurrent variants. The number of affected interactions per hotspot/variant was modeled with a negative binomial. **c**, Average edge betweenness of interactions affected by hotspots and non-recurrent variants. Edge betweenness of an interaction was calculated as the sum of the fraction of all-pairs shortest paths that pass through that interaction in the interactome network (Methods).
